# Beneficial contribution of iPSC-progeny to connexin 47 dynamics during demyelination-remyelination

**DOI:** 10.1101/2020.07.22.216598

**Authors:** Sabah Mozafari, Cyrille Deboux, Cecilia Laterza, Marc Ehrlich, Tanja Kuhlmann, Gianvito Martino, Anne Baron-Van Evercooren

**Author notes:** Corresponding author: Sabah Mozafari Institut du Cerveau et de la Moelle, 47 Bv de l’Hôpital, Paris 75013, France. C. L. present address is: University of Padova, Industrial Engineering Department, Padova, Italy.

## Abstract

Oligodendrocytes are extensively coupled to astrocytes, a phenomenon ensuring glial homeostasis and maintenance of CNS myelin. Molecular disruption of this communication occurs in demyelinating diseases such as multiple sclerosis. Less is known about the vulnerability and reconstruction of the panglial network during adult demyelination-remyelination. Here, we took advantage of LPC-induced demyelination to investigate the expression dynamics of the oligodendrocyte specific connexin 47 (Cx47) and whether this dynamic could be modulated by grafted iPSC-neural progeny. Our data show that deconstruction of the panglial network following demyelination is larger in size than demyelination. Loss of Cx47 expression is timely rescued during remyelination and accelerated by the grafted neural precursors. Moreover, mouse and human iPS-derived oligodendrocytes express Cx47, which co-labels with astrocyte Cx43, indicating their integration into the panglial network. These data suggest that full lesion repair following transplantation occurs by panglial reconstruction in addition to remyelination. Targeting panglial elements by cell therapy or pharmacological compounds may help accelerating or stabilizing re/myelination in myelin disorders.

## Introduction

In the central nervous system (CNS), remyelination of adult axons provides trophic support, rescues loss of function and protect neurons from subsequent axonal degeneration. While remyelination can be provided by endogenous or exogenous progenitors, stem cells, and especially induced pluripotent stem cell (iPSC) derived neural precursors (NPCs) or oligodendrocyte progenitors (OPCs) have become attractive candidates to promote CNS remyelination (Chanoumidou et al., 2020). In particular, intra-parenchymal engraftment of mouse and human iPSCs-NPCs or -OPCs in adult demyelinating conditions results in functional remyelination of host axons (Ehrlich et al., 2017; Mozafari et al., 2015). Grafted NPCs may also provide neuroprotection via trophic support and/or immunomodulation (Ottoboni et al., 2020; Pluchino et al., 2020). As far as cell replacement is concerned, integration of the grafted cells has been evaluated essentially based on their ability to interact with host axons and reconstruct myelin (axo-glial interactions), but their capacity to interact with other glial cells (glial-glial interactions) in order to participate in the pan glial network has been overlooked. This is of major importance since glial-glial interactions are essential for myelin maintenance (Tress et al., 2012) and full repair. Moreover, less is known about how to protect or accelerate panglial repair following adult demyelination.

Astrocytes play an important role in facilitating myelination and myelin maintenance by clearance of extracellular ions and neurotransmitters and by secretion of nutrients including pro-myelinating factors in the CNS (Claycomb et al., 2013; Li et al., 2014). Thus, oligodendrocytes are dependent upon astrocytes for their vital functions and this is primarily controlled by gap junction (GJ) connections. Astrocyte-oligodendrocyte coupling via GJs is crucial for both myelin formation and maintenance, due to K^+^ buffering, energy, and metabolic support for oligodendrocytes via the panglia syncytium (Basu & Sarma, 2018).

In the CNS, the three macroglial cell types, astrocytes, oligodendrocytes, and ependymocytes, are extensively interconnected by GJs channels and form a panglial syncytium. Each of these cells express a set of cell-specific connexins (Cxs) that can connect to make autologous or heterologous GJs (Orthmann-Murphy et al., 2008). Oligodendrocytes specifically express Cx47, Cx45, Cx32 and Cx29, while astrocytes express mainly Cx43, Cx30 and Cx26. Oligodendrocytes, as part of the glial syncytium, are extensively coupled to astrocytes through heterologous GJs such as Cx32/Cx30 and Cx47/Cx43. Cx47/Cx43 channels are largely localized to the oligodendrocyte somata, where they outnumber Cx32/Cx30 channels (Altevogt & Paul, 2004; Kamasawa et al., 2005; Kleopa et al., 2004; Nagy et al., 2003).

Several *in vitro* and *in vivo* studies highlighted that Cx43/Cx47 channels are of major importance for astrocyte/oligodendrocyte crosstalk. It has been proposed that depletion of either Cx43 or Cx47 affects the maintenance of white matter function via alterations in astrocyte-oligodendrocyte homeostasis and play a key role in chronic expansion of demyelinated plaques (Basu & Sarma, 2018). Molecular disruption of astrocytes/oligodendrocytes communication (loss of Cx43/Cx47 connectivity) occurs in demyelinating diseases such as multiple sclerosis (MS) (Markoullis et al., 2014; Masaki, 2015; Masaki et al., 2013; Rash, 2010). Loss of Cx43 in astrocytes is directly correlated with the extent of demyelination and severity of disease in MS and Neuromyelitis Optica (NMO), and extensive loss of Cx43 in the lesion is related to frequent of relapses and rapid disease course (Masaki et al., 2013). While immunoreactivity for astrocyte Cx43 is completely lost within active MS and NMO, it is up-regulated in chronic lesions (Masaki et al., 2013). A significant reduction of astrocyte Cx43, specifically in monocyte infiltrated areas, and a reduction of oligodendrocyte Cx47 in and around the demyelinated plaques occur also in EAE (Brand-Schieber et al., 2005). However, the mechanism of GJ expression alteration in oligodendrocytes has not investigated. In fact, whether demyelinating diseases such as MS with repeated episodes of demyelination could be a cause or a consequence of panglial network deconstruction is not clear.

Genetic alterations of oligodendrocyte or astrocyte connexins lead to myelin disorders. Mutations in the GJC2 gene encoding for human Cx47 cause the hypomyelinating leukodystrophy Pelizaeus–Merzbacher-like disease, whereas mutations in the GJB1 gene encoding Cx32 result in the demyelinating neuropathy X-linked Charcot-Marie-Tooth disease, which can be associated with CNS abnormalities (Kleopa & Scherer, 2006; Uhlenberg et al., 2004). Cx47 ablation in rodents can significantly abolish oligodendrocytes-astrocytes coupling, with the number of coupled oligodendrocytes being reduced by 80 % (Maglione et al., 2010), suggesting a major role for Cx-mediated GJs in oligodendrocyte homeostasis and developmental myelination. Moreover, Cx47 ablation results in myelination abnormalities in and around the compact myelin or periaxonal space and conspicuous vacuolation of nerve fibers in white matter regions. These pathological features are worsened by double deletion of Cx32 and Cx47 resulting in severe myelin alterations and death of the animals (Odermatt et al., 2003). MHV-A59 infection leads to a small, but significant, and persistent downregulation of Cx47 in the MHV-A59 infected brain, and persistent alteration of Cx47 is associated with the loss of the myelin marker PLP in the major white matter tracts of brain (Basu et al., 2017). It has been hypothesized that Cx47 gene mutations affects the permeability of Cx47 channels, which in turn alters exchange of substances between oligodendrocytes or astrocytes, and the way that glia support myelin function (Fasciani et al., 2018). Finally, oligodendrocyte-specific conditional ablation of Cx47 exacerbates acute and chronic relapsing experimental autoimmune encephalomyelitis (EAE) (Zhao et al., 2020). These evidences suggest that oligodendrocyte Cx47 or its astrocyte Cx43 counterpart play an important role in remyelination or myelin pathology. So far only one study reported the dynamic expression pattern of Cx47 in a cuprizone-induced demyelination mouse model (Parenti et al., 2010). However, how oligodendrocytes lose and gain Cx47 expression and whether the panglial repair can be accelerated was not addressed.

We reported that following lysolecithin (LPC)-induced adult demyelination, grafted iPSC-NPCs/OPCs timely differentiate into oligodendrocytes (but also ~ 20% to astrocytes), and extensively remyelinate the lesion 6 weeks after engraftment (Ehrlich et al., 2017; Mozafari et al., 2015). However, the timing and dynamics of loss and gain of Cx47 expression during demyelination and remyelination, and the potential interaction of the iPS-derived oligodendrocytes (through the connexins) with astrocytes during this successful remyelination has not been addressed. Here, we studied in detail the dynamics of Cx47 expression by endogenous and exogenous oligodendrocytes during the demyelination-remyelination process. We find that LPC-induced demyelination causes the loss of the panglial network. While endogenous and exogenous myelinating oligodendrocytes express Cx47 and contribute to the panglial network reconstruction in response to demyelination, exogenous oligodendrocytes accelerate the panglial repair.

## Results

### Oligodendrocytes are frequently connected to astrocytes via Cx43/Cx47 gap junctions in the intact spinal cord

We first questioned how adult murine oligodendrocytes and astrocytes anatomically connect to each other in the spinal cord of adult mice. To this aim, intact spinal cords of adult nude mice (at 8 weeks age) were stained for Cx47 and Cx43, the main oligodendrocyte and astrocyte Cxs respectively. Data show that in the dorsal funiculus, the panglial network consists of CC1^+^ oligodendrocytes expressing Cx47 that were frequently co-labelled with astrocyte Cx43 making a network of glial cells (Fig. 1A). Oligodendrocytes were also connected to each other by homologous Cx47:Cx47, which were distributed mainly on their soma (Fig. 1B, C).

**Figure 1.**
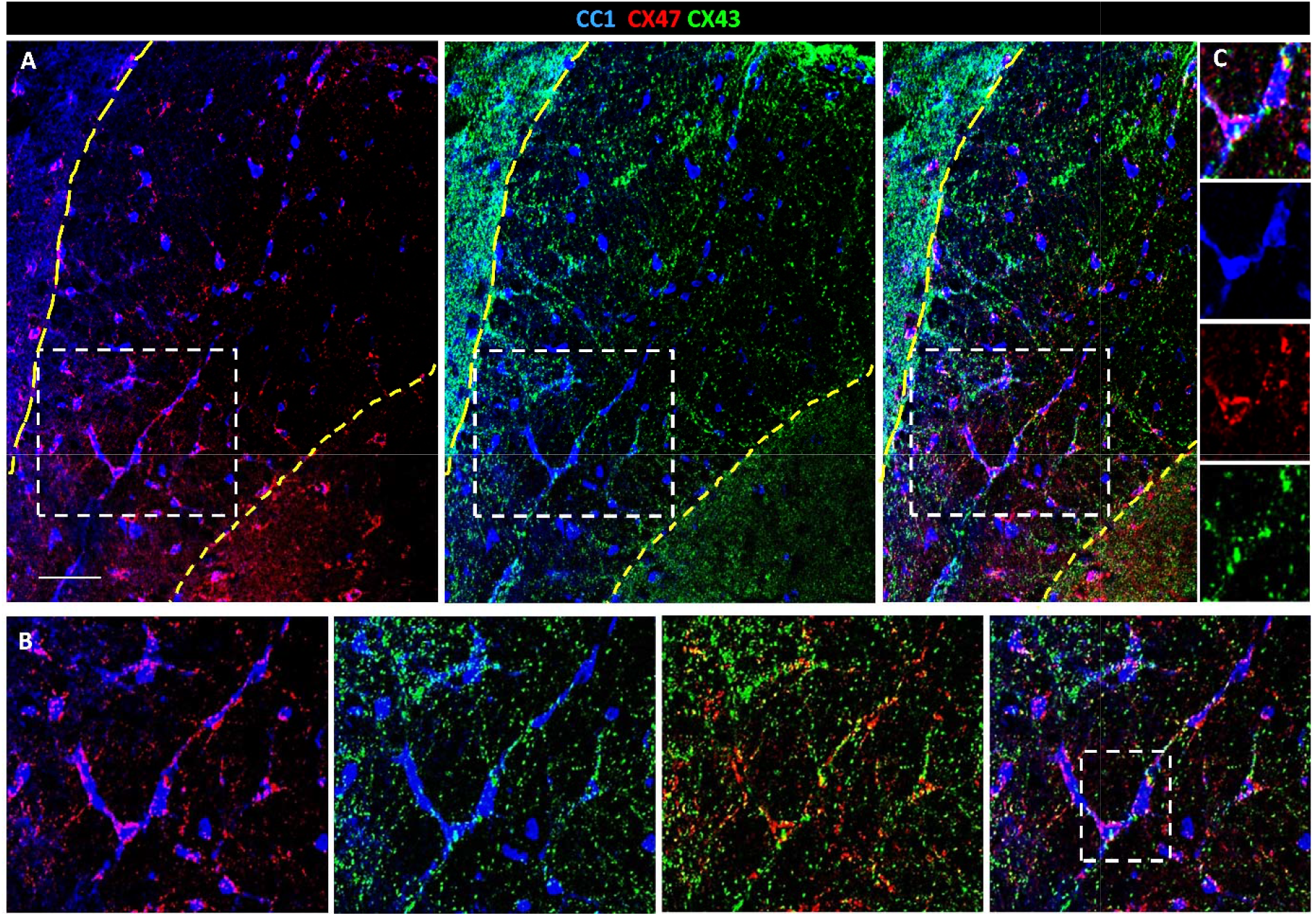
Intact panglial network in the spinal cord of adult nude mice. (A) CC1^+^ (blue) Oligodendrocytes expressing Cx47 (red) together with astrocyte Cx43 (green) make a network of connected glial cells. (B) Higher magnification of the insets in A. (C) Single channels for the inset in B showing two connected oligodendrocytes. Scale bar: 50 μm.

### Deconstruction of the panglial network following LPC-induced demyelination overpasses the extent of demyelination

We first asked what is the consequence of demyelination on the existing pan glial network and whether grafted NPCs would modulate the impact of demyelination. To answer this question, we took advantage of a simplified focal demyelination model injecting 1μl LPC (1%) in the dorsal funiculus of the spinal cord of adult Shiverer:Rag2^−/−^ (myelin-deficient, immune-deficient) or nude (myelin-wild type, immune deficient) mice, (see the study design in Fig. 2A), and injected either medium (1μl) or mouse iPSC- or embryonic brain-derived NPCs (100, 000 cells in 1 μl) in the lesion, 48h after demyelination. The animals were sacrificed 1, 2, 6 and 10 weeks post graft (wpg) or post medium (wpm). Immune deficient animals were used to avoid immune rejection of the transplanted cells and the Shiverer (MBP deficient) mice were used to visualize the graft derived MBP^+^ myelin at later stages.

**Figure 2.**
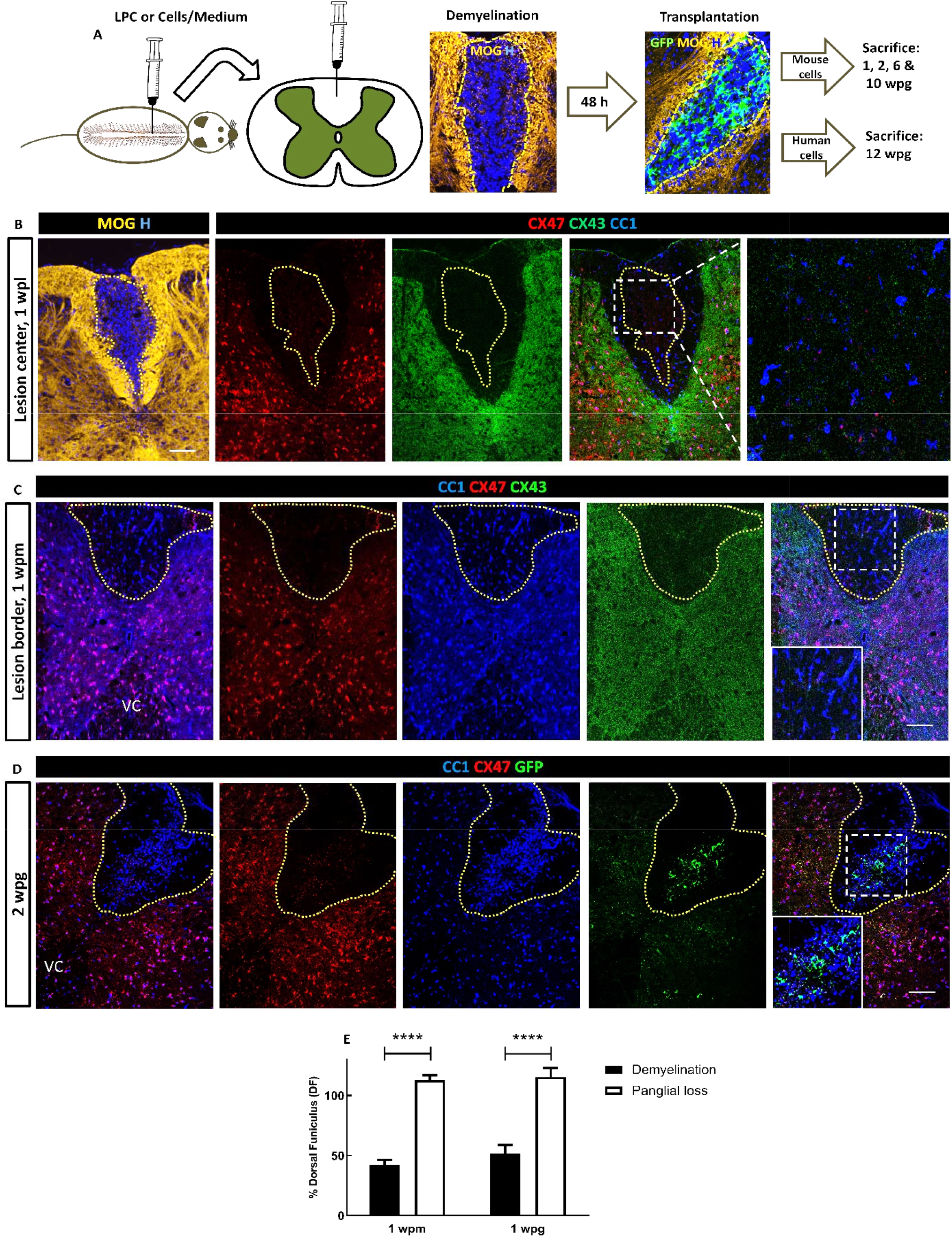
Demyelination disrupts the panglial network beyond the demyelinating zone. (A) Experimental design. Demyelination is induced by LPC injection in the dorsal funiculus (DF) of adult nude or shiverer mice. Mouse iPSC-derived NPCs (miPSC-NPCs), mouse embryonic NPCs (mE-NPCs), or medium were injected 48h after demyelination and animals sacrificed 1 and 2 week post lesion (wpl) in nude mice and 6 or 10 wpl in Shiverer:Rag2^−/−^ mice. A group of shiverer mice were grafted with human iPSC-derived OPCs in the same animal model and sacrificed 12 wpl. (B) At the lesion center, MOG staining (yellow) shows a defined area of LPC induced demyelination at 1 wpl. The loss of Cx47 (red) or Cx43 (green) expression by the CC1^+^ oligodendrocytes (blue) is not limited to the demyelinated area but extends beyond. (C) Although CC1^+^ cells (blue) are present at the lesion border, expression of both oligodendrocyte Cx47 (red) and astrocyte Cx43 (green) is lost while the panglial network at the ventral column (VC) is spared with many CC1^+^/Cx47^+^ or Cx43^+^ cells compared to the dorsal part. (D) The panglial loss following LPC demyelination is still present in a large area around the lesion 2 wpg. (E) Quantifications show that in both medium and cell grafted dorsal funiculi, the panglial loss area (dashed yellow line in C and D) is significantly larger in size than the lesion area (dashed yellow line in B). Two-way ANOVA followed by Tukey’s multiple comparison tests were used for the statistical analysis of these experiments (n=3–6 mice per group). Error bars represent SEMs. ****P < 0.0001. Scale bars: 100 μm.

Analysis of the medium injected mice showed that LPC-induced demyelination disrupted the panglial network as evidenced by the lack of expression of Cx47 and the still obvious reduction of Cx43 expression 1wpm (Fig. 2B). Evaluation of the demyelinated MOG^−^ area and Cx47^−^/Cx43^−^ area at the lesion center revealed that the panglial network disruption was significantly larger in size than the demyelination area (Fig. 2E) with a lack of expression of Cx47 and Cx43 at the lesion border despite the presence of endogenous CC1^+^ oligodendrocytes, at 1wpm compared with non-demyelinated ventral column on the same section containing CC1^+^/C47^+^ cells expressing Cx43^+^ staining, Fig. 2C). The panglial loss was also obvious in grafted lesions and was still present at 2 wpg (Fig. 2D).

### Oligodendrocytes gain Cx47 expression as they repopulate the demyelinated lesion

To study the dynamics of loss and gain of Cx47 expression following demyelination, we quantified the number of CC1^+^/Cx47^+^ oligodendrocytes at the lesion site in nude mice grafted with either miPS-NPCs or mE-NPCs or injected with medium 1 and 2 wpg/wpm. Since no difference in Cx47 expression over time was observed between the two grafted cell types (Fig. S1), data from the two cell types were pooled and compared with those of medium-injected ones. It is of note that our previous comprehensive study comparing side-by-side the two cell types over time in the same animal models showed no significant differences in terms of initial cell density and subsequent proliferation, migration, and differentiation in oligodendrocytes and myelin-forming cells (Mozafari et al., 2015).

Quantification showed that the total number of mature CC1^+^ oligodendrocytes, was significantly increased at 2 wpm/wpg as compared to 1 wpm/wpg with no significant difference between graft vs medium injected groups at both time points (Fig. 3A-E). The percentage of CC1^+^/Cx47^+^ over the total CC1^+^ oligodendrocyte population increased significantly at 2 wpm or 2wpg (Fig. 3F). Almost all of the CC1^+^/Cx47^+^ cells were found at the border of the lesion at 2 weeks post demyelination for both medium and graft injected mice (Fig. 3C and D) with no CC1^+^/C47^+^ oligodendrocytes at the lesion core at 1 and 2 wpm/wpg (see insets in Fig. 3A-D). In the grafted mice, only few GFP^+^/CC1^+^/Cx47^+^ cells were found with small amounts of Cx47 expression and no significant increase at 2 vs 1 wpg (Fig. 3G).

**Figure 3.**
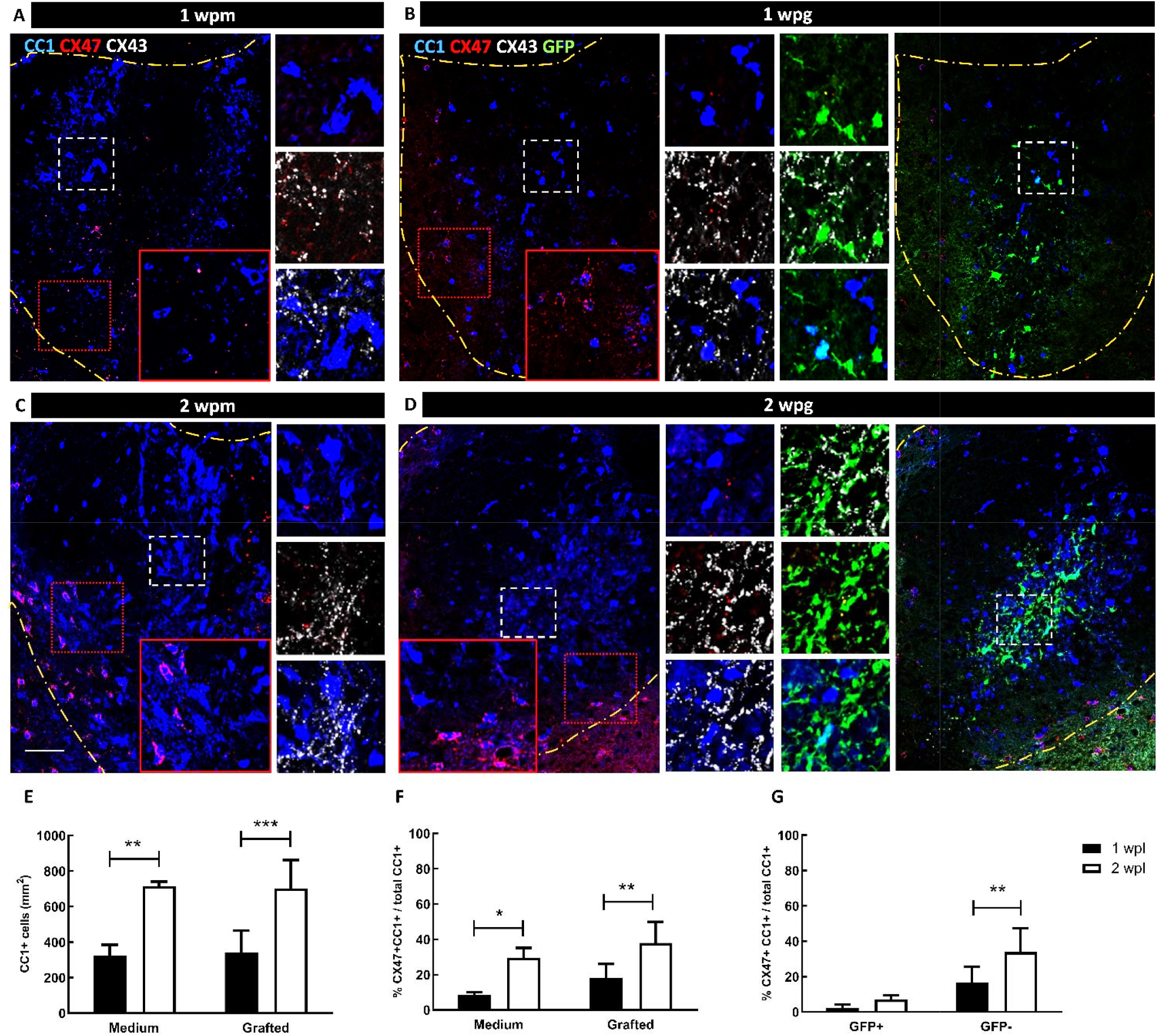
Dynamics of oligodendrocyte-Cx47 loss and gain of expression by endogenous and exogenous cells following demyelination. (A and B) One wpl in medium and cell injected nude mice spinal cords, the number of CC1^+^ cells (blue) was decreased, and both endogenous or grafted GFP^+^ cells (green) were almost entirely negative for the oligodendrocyte-specific Cx47 marker (red) at the center of the lesion (inset boxes with dashed white lines). Few oligodendrocytes were co-labelled with few astrocyte specific Cx43 marker (white). (C-E) The number of CC1^+^ cells increased significantly 2 wpl with a slight increase in CC1^+^/Cx47^+^ (found mainly at the border f dorsal funiculus, red inset boxes) as compared to the first wpl. Dashed yellow lines show the limits of dorsal funiculus. (F) Quantification indicates that the percentage of Cx47^+^/CC1^+^ cells was significantly increased at 2 wpl in both medium and grafted animals. (G) In the grafted animals, the percentage of GFP^+^/Cx47^+^ oligodendrocytes did not increase significantly over time, while the percentage of Cx47^+^/GFP^−^ endogenous oligodendrocytes did so. Two-way ANOVA followed by Tukey’s multiple comparison tests were used for the statistical analysis of these experiments (n=3–6 mice per group). Error bars represent SEMs. **P < 0.001. wpg: week(s) post graft; wpm: week(s) post medium; wpl: weeks post lesion. Scale bar: 50 μm.

Next, we asked whether the grafted cells had any impact on the percentage of endogenous oligodendrocyte Cx47 expression. The percentage of GFP^−^/CC1^+^/Cx47^+^ over total CC1^+^ cells increased significantly in the grafted mice at 1 and 2 wpg (Fig. 3G) suggesting that the grafted cells induced Cx47 expression by endogenous oligodendrocytes. Interestingly, some CC1+ cells slightly co-labelled with astrocyte Cx43 at 1 wpg and their number increased slightly at 2 wpg (insets in Fig. 3A-D). GFP^+^/Cx43^+^ cells were the source of many of these Cx43^+^ plaques at 1 or 2 wpg consistent with the observation that many GFP^+^ expressed GFAP^+^/Nestin^+^ around lesion 1 wpg (Fig. S2).

### Gain of Cx47 expression during remyelination is accelerated by exogenous NPCs

We previously demonstrated that miPS-derived NPCs competitively remyelinated numerous demyelinated host axons with compact myelin, restored nodes of Ranvier and rescued the delayed conductance velocity of Shi/Shi Rag2^−/−^ mice to the same extent than brain-NPCs (Mozafari et al., 2015). Here we aimed to study in the same context, the effect of the graft on the temporal dynamics of Cx47 expression during the remyelination process. To this end, we quantified the number of the CC1^+/^Cx47^+^ cells in the intact, grafted, and medium injected control mice, at 6 and 10 weeks after spinal cord demyelination (Fig. 4).

**Figure 4.**
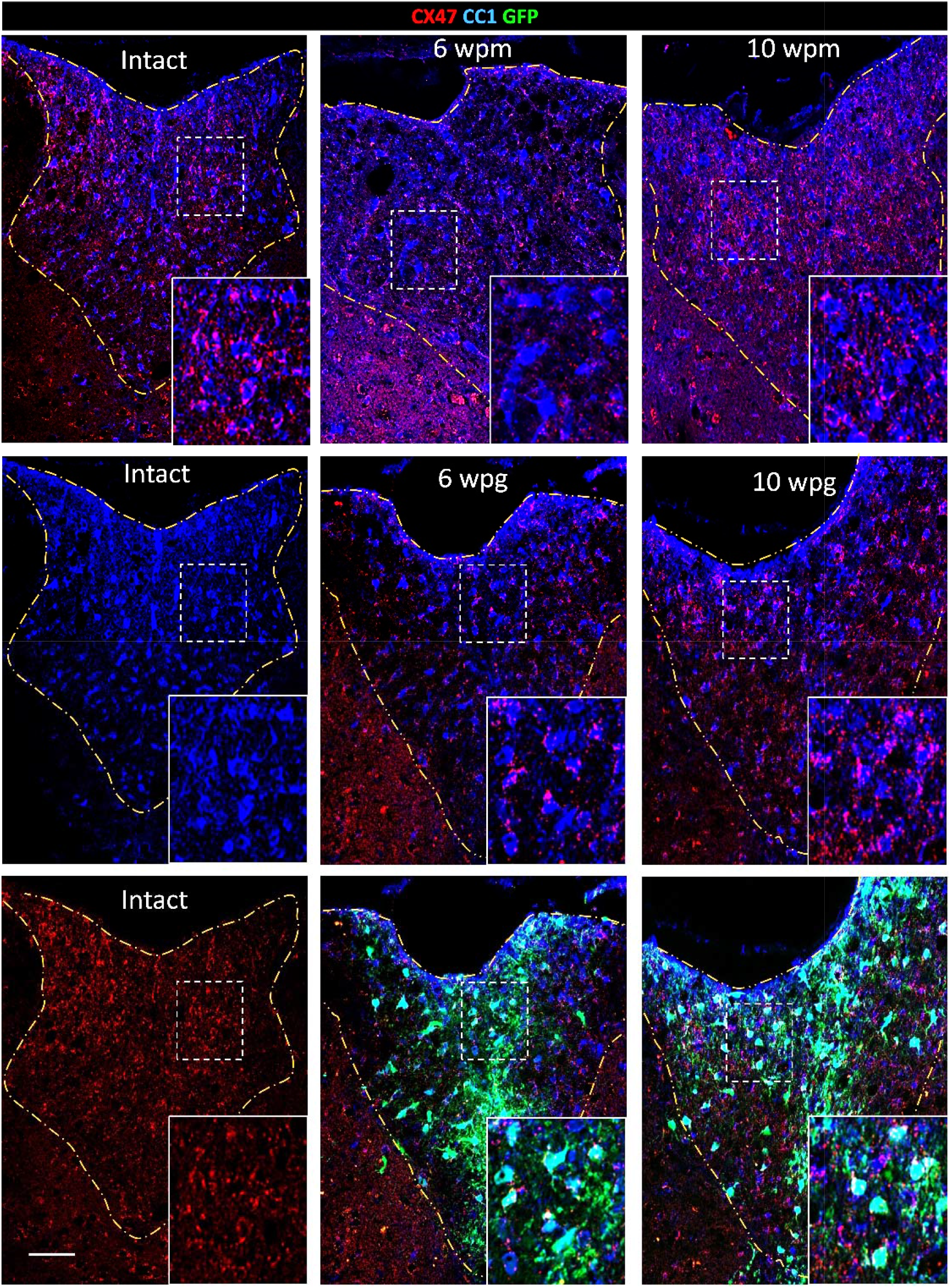
Dynamics of oligodendrocyte Cx47 gain of expression during endogenous and exogenous remyelination. Expression of Cx47 (red) by CC1^+^ oligodendrocytes (blue) at 6 and 10 wpl in the dorsal funiculus (dashed yellow lines) of Shiverer:Rag2^−/−^ mice, followed by medium injection (wpm) or engraftment (wpg) of miPSC-NPCs was compared to a group of intact mice. At 6 wpg, CC1^+^ cells expressed more Cx47^+^ gap junction (GJ) plaques compared to 6 wpm. Yet, there was no difference with intact and 10 wpm mice. At 10 wpg, CC1^+^ oligodendrocytes expressed more Cx47^+^ GJ plaques compared to 6wpg, 10 wpm and intact mice. Scale bar: 50 μm. wpg: week(s) post graft; wpm: week(s) post medium; wpl: weeks post lesion. Quantitative data are reported in Figure 5.

In the medium group, the total number of CC1^+^ cells were reduced at 6 wpm as compared to intact mice but was rescued at 10 wpm (Fig. 5A). The total number of CC1^+^ cells did not increase after engraftment (Fig. 5A). As for the percentage of CC1^+^/Cx47^+^ cells (Fig. 5B), it was significantly lower at 6 wpm as compared with intact but was partially restored at 10wpm to levels of intact mice. In contrast the percentage of CC1^+^/Cx47^+^ cells in the transplanted group remained steady at 6 wpg, and 10 wpg, and was comparable to the levels of intact mice. Fig. 4 showed that the amount of Cx47^+^ GJ varies among the intact, medium and grafted mice. To gain insight on the differences in oligodendrocyte Cx47 expression levels per group, we quantified the number of Cx47 gap junction (GJ) plaques expression per CC1^+^ cells and namely CC1+ cells with 5 or less (≤ 5, Cx47^low^) and those with more than 5 (> 5, Cx47^high^) Cx47 GJ plaques. In the intact shiverer mice, nearly 48% of CC1^+^ cells were Cx47^low^ and 49% were Cx47^high^ (Fig 5. C and D). The percentage of CC1^+^/Cx47^low^ cells was significantly higher in the medium group at 6 wpm (74 %) as compared to the intact, and grafted groups at 6 wpg and 10 wpg. At 10 wpm the percentage of CC1^+^/Cx47^low^ cells (Fig. 5 C) was significantly higher (65 %) than that in the intact and 10 wpg mice. The CC1^+/^Cx47^low^ population represented 59 % and 29 % of total CC1^+^ cells respectively at 6 and 10 wpg. The percentage of CC1^+^/Cx47^high^ cells (Fig. 5D) increased significantly over time for both medium treated and grafted mice (from 12 % at 6 wpm to 28 % at 10 wpm, and 35 % at 6 wpg to 68 % at 10 wpg). Interestingly the higher percentage of CC1^+^/Cx47^high^ cells in the grafted group at 10wpg was significantly higher than that of intact mice. These data suggest that during remyelination in transplanted mice, Cx47 expression levels were not only accelerated compared to medium-treated mice, but also exceeded that of intact mice.

**Figure 5.**
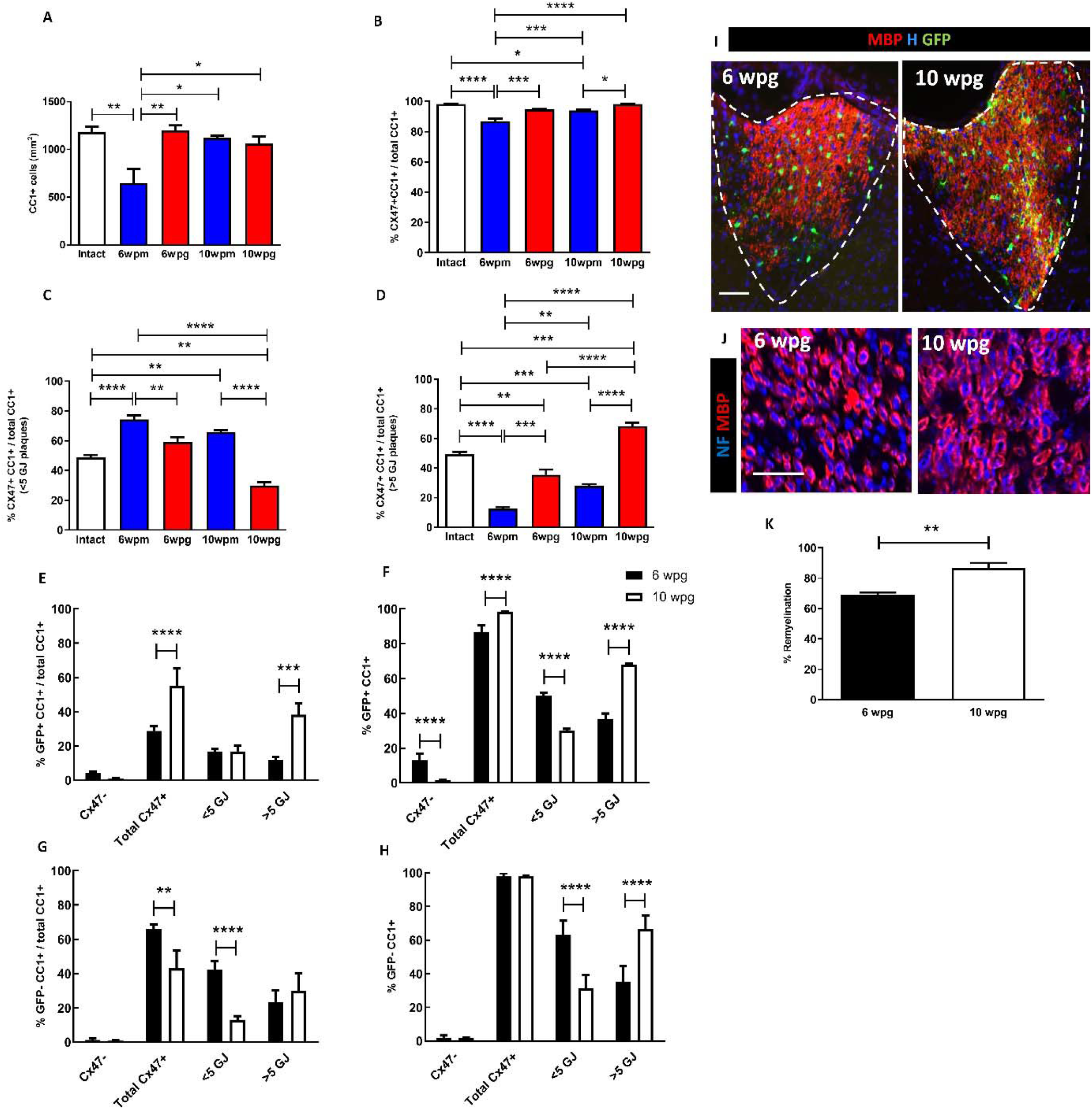
The dynamic expression of Cx47 following demyelination, is accelerated by grafted NPCs. (A) The total number of CC1^+^ cells in the lesion was significantly lower at 6 wpm as compared to intact, 6 wpg, 10 wpm and 10 wpg mice. The total number of CC1^+^ cells did not increase after engraftment. (B) The decrease in CC1^+^/Cx47^+^ cells in 6 wpm mice over intact mice was rescued in 6 wpg mice. While at 10 wpm, the percentage of CC1^+^/Cx47^+^ cells was not fully recovered to that of intact mice, it was fully rescued in 10 wpg mice. No significant difference was observed in the percentage of CC1^+^/Cx47^+^ cells between 6 vs 10 wpg (see figure 4). (C) The percentage of CC1^+^/Cx47^low^ cells in 6 wpm and 10 wpm mice was significantly higher than that in intact mice. CC1^+^ oligodendrocytes expressed lower amounts of Cx47 plaques respectively at 6 wpg and 10 wpg as compared to 6 wpm and 10 wpm. In 10 wpg mice, the percentage of CC1^+^/Cx47^low^ cells was significantly lower than that of intact mice. (D) The percentage of CC1^+^/Cx47^high^ was significantly lower in 6 wpm, 10 wpm and 6 wpg compared to intact mice, but rescued over intact mice at 10 wpg. (E) Grafted cells constitute more than 50% of total CC1^+^/Cx47^+^ cells at 10 wpg with most of them expressing high levels of Cx47 GJ plaques. (F) The expression of Cx47 among the GFP^+^/CC1^+^ population increased significantly over time with higher Cx47 expression at 10 wpg vs 6 wpg. (G) In the grafted mice, endogenous CC1^+^/Cx47^+^ cells constitute a higher proportion of total CC1+ cells (most of them were CC1^+^/Cx47^low^ cells) at 6 wpg but decreased over time (compare with graph E). (H) In the grafted mice, excluding the grafted population, almost 100% of endogenous CC1^+^ cells expressed Cx47 at 6 wpg as well as 10 wpg and gained more Cx47 GJ plaques over time (See Fig. 4). (I) Remyelination (MBP expression in red) of the grafted cells (green) in dorsal funiculus increased over time. (J and K) The percentage of NF^+^ host axons (blue) wrapped with exogenous MBP^+^ myelin increased significantly over time. One-way followed by Tukey’s (graphs in A-D), Two-way ANOVA followed by Sidak’(graphs in F-H) multiple comparison tests or Mann Whitney test (graph in K) were used for the statistical analysis of these experiments (n=3 mice per group). Error bars represent SEMs. * P < 0.05, **P < 0.001, *** or **** P < 0.0001. wpg: week(s) post graft; wpm: week(s) post medium. Scale bars: 50 μm (in I) and 10 μm (in J).

Next, we questioned the relative contributions of grafted (GFP+) and endogenous (GFP−) oligodendrocytes to the Cx47 GJ expression in grafted mice (Fig. 5E and G). Data show that the GFP^+^/CC1^+^/Cx47^+^ oligodendrocytes consist of 28 % and 55% of total CC1^+^ cells respectively at 6 and 10 wpg. In addition, at 10wpg, the GFP^+^/ CC1^+^/Cx47^high^ pool contributed majorly (38%) to the GFP^+^/CC1^+^/Cx47^+^ increase (Fig. 5E). In addition, the percentage of Cx47^+^ cells as well as Cx47 level of expression increased over time in the grafted oligodendrocyte population, (Fig. 5F) with 86% and 98% of GFP^+^/CC1^+^ cells expressing Cx47 respectively at 6 wpg (50% Cx47^low^ and 36% Cx47^high^) and 10 wpg (30% Cx47^low^ and 68% Cx47^high^).

How transplanted cells impact on endogenous Cx47^+^ oligodendrocytes was our next question. Quantification showed that in the transplanted shiverer mice, endogenous GFP^−^/CC1^+^/Cx47^+^ consist of 66% of the total CC1^+^ cells with 42% Cx47^low^ at 6 wpg (Fig. 5G). As the percentage of GFP^+^/CC1^+^/Cx47^+^ cells increased to 55% at 10 wpg, the percentage of endogenous shiverer GFP^−^/CC1^+^/Cx47^+^ cells decreased to 43% taking into account that the total number of CC1+ cells does not fluctuate in the grafted mice. However, 98% of the endogenous population expressed Cx47 with 63% Cx47^low^ cells and 66% Cx47^high^ respectively 6 and 10 wpg (Fig. 5H).

These data altogether show that during lesion repair, the number of Cx47^+^ GJ plaques per CC1^+^ cells increases with time within the total CC1^+^ cells or within the endogenous or GFP^+^ transplanted oligodendrocyte population.

We then asked whether this increase could be correlated to progress of remyelination itself. Immunolabeling for MBP and NF revealed that 69% of the axons in the lesion core were remyelinated by the grafted cells at 6 wpg. The proportion of remyelinated axons increased significantly from 68% at 6wpg up to 86% at 10wpg (Fig. 5I-K). Comparing data from graph F with K (Fig. 5) shows that the increase in the number of Cx47^+^ oligodendrocytes and in the level of Cx47 expression per cell, fluctuate with the progressive increase in remyelination, suggesting that Cx47 expression and remyelination are temporally correlated.

### The iPSC-derived neural progeny integrates in the panglial network after transplantation in dysmyelinated mice

Imaging at high magnification confirm that mature CC1^+^/Olig2^+^ host- and grafted mouse iPSC-oligodendrocytes were able to express Cx47 (Fig. 6A) during remyelination. We also observed that not only mature CC1^+^ oligodendrocytes but also CC1^−^/Olig2^+^ immature oligodendrocyte progenitor cells (empty white arrowheads insets in Fig. 6A) express Cx47 and make homologous gap junction Cx47-CX47 with adjacent oligodendrocytes (Fig. 6A insets).

**Figure 6.**
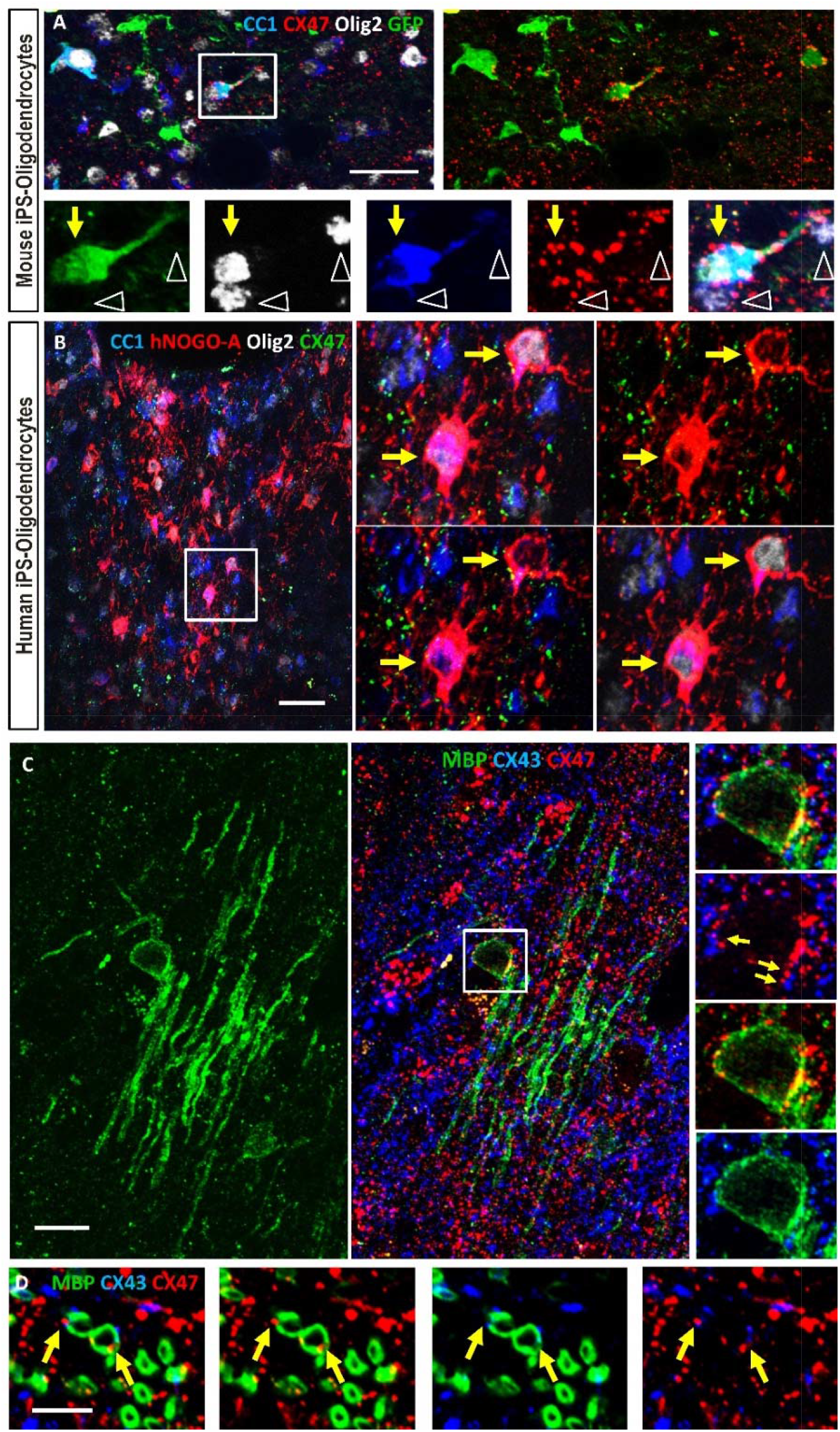
iPSC-derived oligodendrocytes have the molecular machinery to interact with other glial cells. (A) Mouse GFP^+^ (green)/CC1^+^ (blue)/Olig2+ (grey) iPSC-derived oligodendrocyte expressing Cx47 (red). (B) Human iPSC-derived CC1^+^ (blue)/Olig2^+^ (grey)/hNOGOA^+^ (red) oligodendroglia expressing Cx47 (green) integrated into the demyelinated dorsal funiculus. (C) A mouse iPS-derived oligodendrocyte expressing MBP^+^ myelin (green) connected to astrocytes via heterologous GJs Cx47:Cx43 (red:blue) on their soma (inset, yellow arrows). (D) Graft-derived MBP^+^ myelin-like structures seen in cross sections (green) decorated by Cx47^+^ gap junction plaques (red). Scale bars: 50 μm (in A and B), 25 μm (in C) and 5 μm (in D).

**Figure 7.**
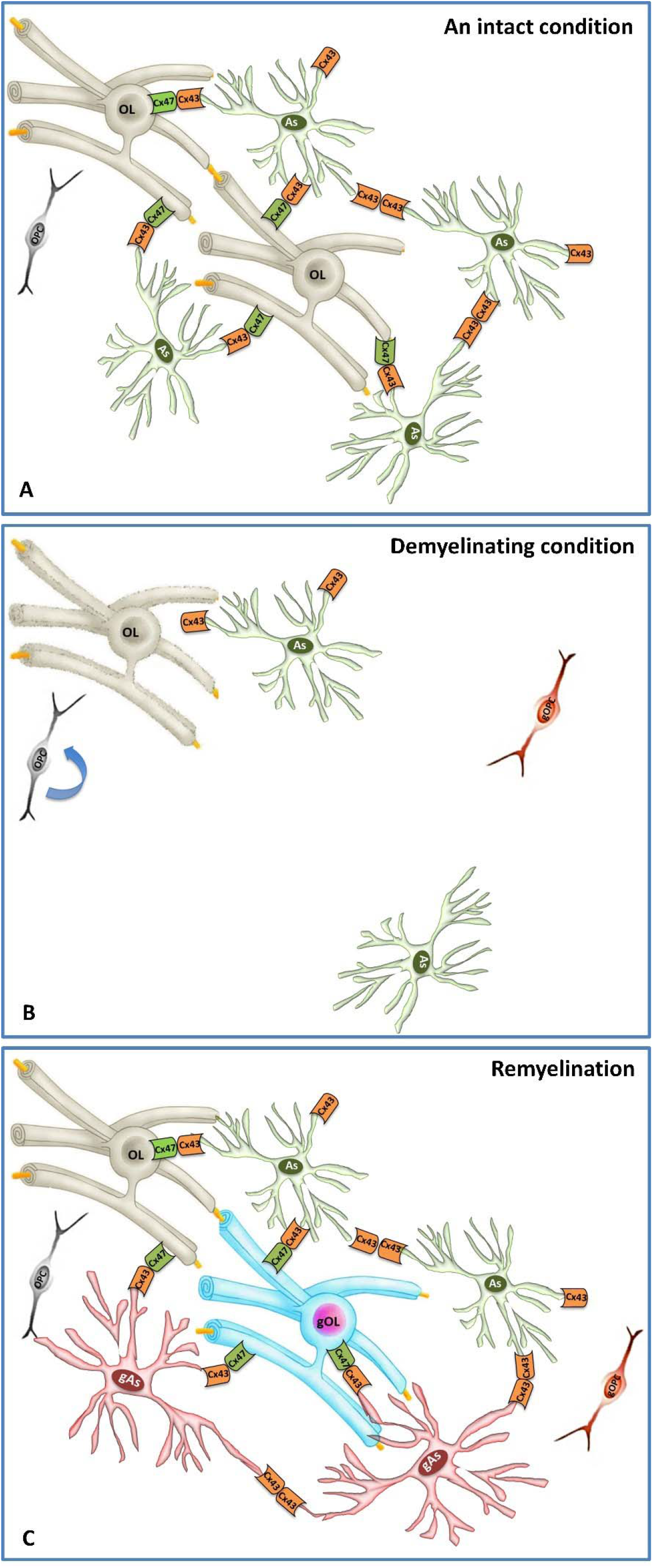
Graphical abstract of the oligodendrocyte-astrocyte syncytium dynamics in response to demyelination. (A) The adult glial syncytium consists of astrocytes and oligodendrocytes interacting mainly through the heterologous gap junction complex Cx43:Cx47 (showed in orange:green boxes). (B) During demyelination, this network is largely deconstructed. (C) During reyelination, the glial syncytium is reconstructed which could be accelerated by graft-derived remyelination. OL: oligodendrocyte; As: astrocyte; OPC: oligodendrocyte precursor cell; gOL: graft-derived oligodendrocyte; gAs: graft-derived astrocyte; gOPC: grafted OPC.

To investigate whether human iPSC-derived oligodendrocytes were able to express Cx47 following demyelination, human iPSC-derived O4^+^ cells (hiOL) were also grafted in the adult demyelinated spinal cord of the Shiverer:Rag2^−/−^ mice (see Fig. 2A) and animals sacrificed 12 wpg. In these conditions, hiOL vastly remyelinate the demyelinated spinal cord (Ehrlich et al., 2017). Here we show that the human iPSC-derived oligodendrocytes identified by CC1, human NOGOA and Olig2 expressed also Cx47 (Fig. 6B). Moreover, the iPSC-derived oligodendrocytes were frequently co-labelled with astrocyte Cx43 to make Cx47^+^/Cx43^+^ heterologous gap junctions (Fig. 6C). Cx47^+^/Cx43^+^ co-labeling was found not only on iPSC-oligodendrocyte somas but also on iPSC-derived myelin (Fig. 6D). These data suggest that rodent and human iPS-derived oligodendrocytes have the molecular machinery to interact with other glial cells during remyelination and participate in reconstructing the panglial network lost to demyelination.

## Discussion

Alterations in connexin’s expression by oligodendrocytes or associated astrocytes suggest that gap junction components that connect one cell to another play a major role in genetic or acquired myelin disorders (Cotrina & Nedergaard, 2012; Xia et al., 2020). While most studies related to myelin diseases have focused on how to increase re/myelination via axo-glia interactions as a mean to protect and insure axon’s proper function, less is known about the panglial syncytium reconstruction in order to provide maintenance of the newly generated myelin and thereby secure full repair of the lesion. We have used LPC-induced demyelination to investigate in detail the dynamic expression of oligodendrocyte Cx47 during the demyelination-remyelination process of the adult murine spinal cord. Moreover, we grafted iPSc-NPCs/OPCs following LPC demyelination to explore whether they play role in the loss and reconstruction of the glial-glial network.

We show that in the intact adult spinal cord of myelin-wild type nude mice, oligodendrocytes express Cx47 and connect to one another, as well as to astrocytes via Cx47 and Cx43 expression as previously reported in the intact mouse corpus callosum (Maglione et al., 2010). LPC-induced focal demyelination alters the panglial network and more specifically disrupts Cx47-Cx47 and Cx47-Cx43 connections. Interestingly, this loss extends beyond the size of demyelination suggesting that the white matter panglial network is highly sensitive to demyelination, and that a given area of myelin is supported by a larger area of glial-glial connections. This is in line with observations arising from chronic MS lesions both in white and gray matter as well as in normal appearing white matter (NAWM) (Markoullis, Sargiannidou, Schiza, et al., 2012; Markoullis et al., 2014). In MS, Cx loss is correlated with inflammation and disease duration, suggesting that oligodendrocyte disconnection from reactive astrocytes may play a role in failed remyelination and disease progression (Markoullis, Sargiannidou, Schiza, et al., 2012). The confirmation of these observations in the MS animal model, EAE, suggest that GJ deficient myelinated fibers appear more vulnerable to CNS inflammatory demyelination, and that persistent impairment of both intra- and intercellular oligodendrocyte GJs even in NAWM, may be an important mechanism of MS progression (Markoullis, Sargiannidou, Gardner, et al., 2012). The present data imply that LPC-induced focal demyelination might provide a valuable simplified model to study adult panglial loss observed in demyelinating diseases.

Our data revealed that 2 weeks following demyelination, only few oligodendrocytes around the lesion start to re-express Cx47 despite the increasing number of oligodendrocytes at the lesion site. Down and up-regulation of Cx47 occur also in cuprizone-induced demyelination (Parenti et al., 2010) and EAE (Markoullis et al., 2012a). In the later, Cx47 expression increased during remyelination at 28 dpi but decreased again at 50 dpi in the relapsing phase (Markoullis, Sargiannidou, Gardner, et al., 2012). These observations indicate that Cx expression is a very sensitive and dynamic process occurring in response to any type of demyelination.

We also provide evidence that, engraftment of mouse embryonic or iPSC-derived NPCs results in a higher percentage of endogenous oligodendrocytes expressing Cx47. Interestingly, one study revealed that intraventricular transplantation of NPCs 7 days following EAE induction prevents the reduction of oligodendrocyte Cx47, and increased the number of oligodendrocytes, local Cx47 levels and Cx47 GJ plaques per cell suggesting a neuroprotective role of the transplanted NPCs on the panglial network (Theotokis et al., 2015). How exogenous NPCs induce Cx47 expression in endogenous oligodendrocytes is not known. Yet, an increasing body of evidences support that NPC’s neuroprotective effect on endogenous cells occurs via cytokines and/or neurotrophic factors (Laterza et al., 2013; Marteyn et al., 2016; Ottoboni et al., 2020; Pluchino et al., 2020). Moreover, we show that the grafted cells differentiate into oligodendrocytes as well as in GFAP+ astrocytes, which in turn, could help promoting Cx47 expression in endogenous oligodendrocytes (May et al., 2013). Jaderstad et al. showed that grafted NPCs into a cerebellar organotypic slice culture system, have a neuroprotective effect on host neurons through the gap junction they make with neurons particularly mediated by astrocytic Cx43 (Jaderstad et al., 2010). Another study shows that ciliary neurotrophic factor (CNTF) activates the JAK/STAT pathway leading to enhanced Cx43 expression and intercellular coupling (Ozog et al., 2004). Whether Cx47 (or its major partner Cx43) can be upregulated by trophic factors secreted from NPCs needs to be investigated.

We found that the percentage of oligodendrocytes expressing Cx47 is accelerated in the grafted animals 6 and 10 wpg compared to medium-injected groups. Moreover, the percentage of oligodendrocytes expressing higher levels of Cx47 increases significantly over time in the grafted animals. Endogenous oligodendrocytes constitute a higher percentage of Cx47+ cells with most of them expressing low levels of Cx47 GJ plaques at 6 wpg. However, at 10 wpg, grafted cells harbor a greater proportion of the total Cx47+ oligodendrocyte population, with most of them expressing high levels of Cx47 GJ plaques, thus suggesting a higher contribution of graft-derived oligodendrocytes in total Cx47 expression over time. Moreover, the percentage of endogenous oligodendrocytes with high levels of Cx47 is clearly higher in the grafted vs medium groups (28 % at 10 wpm vs 66 % at 10 wpg) indicating, once more, the beneficial effect of grafted cells on the endogenous cells. In this context, future studies using loss of function experiments in which expression of Cx47 in the grafted cells is blocked should provide valuable information.

We previously demonstrated that following transplantation of mouse or human iPSC-derived NPCs or OPCs in demyelinated spinal cord of Shi/Shi Rag2^−/−^ mice, they extensively remyelinate the lesion (Ehrlich et al., 2017; Mozafari et al., 2015). Here we show that the progressive increase in oligodendrocyte Cx47 expression is timely correlated with the increase in remyelination by the grafted cells. These data altogether, suggest that the more oligodendrocytes produce myelin around axons the more they express Cx47 GJ plaques, highlighting the need of highly remyelinating cells to firmly integrate into the glial network to receive necessary metabolic/homeostasis supports.

We also show that grafted cells express oligodendrocyte Cx47 which frequently co-labels with astrocyte Cx43 during remyelination. The observation that exogenous and endogenous oligodendrocytes co-express oligodendrocyte Cx47 and astrocyte Cx43 is in line with the fact that NPC engraftment leads to reactivity of endogenous astrocytes as well as the differentiation of a small population of the grafted cells into astrocytes, indicating that both astrocyte sources could contribute to the astrocyte Cx43 expression on oligodendrocyte. Thus, mouse and human skin-derived remyelinating oligodendrocytes are capable of being integrated into the panglial network by establishing proper molecular connections with other glial cells to ensure their function. It is known that Cx43-Cx47 co-expression enables astrocytes to fuel oligodendrocyte the necessary energy supply through gap junction GJ-mediated communication (Giaume et al., 2013; Hirrlinger & Nave, 2014). A mutated form of Cx43, which is only delivered to plasma membrane, but does not form functional channel, shows that the presence of Cx43 at the cell surface, is necessary and sufficient for normal expression, phosphorylation and stability of Cx47-mediated GJ plaques at the cell surface (independent of Cx43 GJC function on the cell surface), suggesting a dependency of oligodendrocyte Cx47 GJ channel stability on astrocyte Cx43 expression (May et al., 2013).

Although our data clearly support the dynamic expression of Cx47 together with Cx43 during the demyelination-remyelination process, many open questions remain to be answered, including the mechanism involved in the endogenous-exogenous interface during remyelination which could lead to the means to pharmacologically accelerate panglial repair following demyelination, the capacity of the panglial network to help support myelin in repeated episodes of dem/remyelination, the functionality of the established connections and finally the respective roles of astrocyte or oligodendrocyte Cxs in lesion expansion, and OPCs differentiation and remyelination in order to better clarify the functional consequences of these networks.

Altogether, the data of the present study provide valuable findings in the biology and dynamic oligodendrocyte Cx47 expression during adult demyelination and remyelination. More specifically we show that transplantation of NPCs might help improving remyelination of the lesion, but also participate in and accelerate reconstruction of the panglial network to ensure their functional integration and maintenance of newly generated myelin. Data from this study along with others, indicate that Cx-mediated communication among oligodendrocytes, among astrocytes, or between astrocytes and oligodendrocytes may be important for oligodendrocyte development during re/myelination. Despite the lack of definite evidence for the role of GJs in the pathology of adult demyelinating diseases such as MS, these findings shed light on the mechanisms involved in myelin pathology and repair *in vivo* and can imply the role of GJs as contributors or modifying factors during MS pathogenesis or therapy. Deciphering the factors involved in Cx-mediated regulation of oligodendrocyte differentiation or remyelination may provide clues for novel strategies to rescue hereditary dysmyelinating diseases and improve remyelination by manipulating Cx expression or enhancing GJ channel function in acquired demyelinating diseases.

## Materials and methods

### Cell culture

#### Mouse iPS-NPCs

miPSCs were generated from reprogrammed E14.5 C57Bl/6 *Sox2^ßgeo/ßgeo^* knock-in embryonic fibroblasts expressing a ßgeo cassette under the control of Sox2 promoter that provides cell resistance to neomycin (Laterza et al., 2013). NPCs were grown on uncoated plastic flasks (T75) in Euromed-N medium (Euroclone) supplemented with 1% N2 nutrients (Invitrogen), Pen/strep (Invitrogen), L-glutamine (200nM) (Invitrogen) and 20ng/ml basic fibroblast growth factor (bFGF) (Preprotech) and 20ng/ml epidermal growth factor (EGF) (Preprotech). Fresh medium was added every other day. Cells were split once a week depending on cell density and were reseeded at a density of 10^6^ cells/T75 flask in 10 ml of medium.

#### Mouse embryonic NPCs (mE-NPCs)

Primary mouse NPCs were obtained from E12/E13 C57Bl/6 mice as previously described (Deboux et al., 2013). Briefly, brains were dissected free of meninges, dissociated using ATV (0.05% trypsin, 0.1% glucose and 0.5 nM EDTA). Collected cells were resuspended in NEF medium composed of DMEM/F12 medium (1:1) supplemented with N2 nutrients (1%), B27 (0.5%), insulin (25 μg/ml), glucose (6 mg/ml), Hepes (5 mM), FGF2 (20 ng/ml) and EGF (20 ng/ml). NPCs were dissociated once a week and reseeded at the density of 10^6^ cells/T75 flask in 10 ml medium. Immunocytochemistry showed that miPS-NPCs similar to their embryonic counterparts expressed essentially Olig2 (97 %), nestin (98 %), and Ki67 (71 %) while they were negative for OPC markers PDGFRα and O4 or neuronal marker MAP2 *in vitro*, before engraftment (Mozafari et al., 2015). For *in vivo* cell tracking, NPCs were transduced with a third-generation lentiviral vector ancoding the green fluorescent protein GFP. More than 80% of the cells were labeled with this method (Laterza et al., 2013).

#### Human iPSC-OPCs

The methods to generate and maintain the human OPCs from iPSCs were previously described (Ehrlich et al., 2017). Briefly, iPSCs were differentiated into NPCs by treatment with small molecules as described (Ehrlich et al., 2017; Reinhardt et al., 2013). Differentiation of NPCs into O4+ oligodendroglial cells was achieved with a poly-cistronic lentiviral vector containing the coding regions of the human transcription factors SOX10, OLIG2 and NKX6.2 (SON) followed by an IRES-pac cassette allowing puromycin selection for 16h (Ehrlich et al., 2017). Human NPCs were seeded at 1.5 × 10^5^ cells/well in 12-well plates, allowed to attach overnight and transduced with SON lentiviral particles and 5 μg/ml protamine sulfate in fresh NPC medium. After extensive washing, viral medium was replaced with glial induction medium (GIM). GIM was replaced after 4 days with differentiation medium (DM). After 12 days of differentiation, cells were dissociated by accutase treatment for 10 min at 37°C, washed with PBS, re-suspended in PBS/0.5% BSA buffer and singularized cells filtered through a 70 μm cell strainer (BD Falcon). Cells were incubated with mouse IgM anti-O4-APC antibody (Miltenyi Biotech) following the manufacturer’s protocol, washed, re-suspended in PBS/0.5% BSA buffer (5 × 10^6^ cells/ml) and purified by magnetic cell sorting using anti-O4 MicroBeads (Miltenyi Biotec) following the manufacturer’s protocol. Media details were provided in (Ehrlich et al., 2017).

*In vitro* characterization by immunocytochemistry revealed that the human iPS-OPCs were NG2+ and highly expressed GALC and O4 (70 %) 28 days *in vitro* (Ehrlich et al., 2017). MACS-purified O4+ cells at 14 days *in vitro*, were used for *in vivo* studies.

### Animals

To study the dynamic expression of oligodendrocyte Cx47 following demyelination and remyelination after engraftment, we used two different immunodeficient mouse strains: nude mice with normal myelination and *Shi/Shi:Rag2^−/−^* with dysmyelination (MBP deficient mice) backgrounds as previously published (Mozafari et al., 2015). *Nude mice* were adult *RJ:NMR-Fonx1nu/Fonx1nu* immunodeficient mice (n=20, 8-9 weeks of age, Janvier). *Shiverer* mice were crossed to *Rag2* null immunodeficient mice (Shinkai et al., 1992) to generate a line of *Shi/Shi:Rag2^−/−^* dysmyelinating immunodeficient mice (n=21, 8-9 weeks of age). Mice were housed under standard conditions of 12-hours light/12 hour dark with ad libitum access to dry food and water cycle at ICM institute’s animal facility. Experiments were performed according to European Community regulations and were approved by the National Ethic’s Committee (authorization 75-348; 20/04/2005) and local Darwin Ethic’s Committee.

### Demyelination and cell transplantation

To induce demyelination, mice were anaesthetized by intraperitoneal injection of a mixture of 100 mg/kg Ketamine (Alcyon) and 10 mg/kg Xylazine (Alcyon). Focal demyelination was performed as previously described (Blanchard et al., 2013; Buchet et al., 2011; Mozafari et al., 2015) by stereotaxic injection of 1μl of 1% lysolecithin (LPC; Sigma-Aldrich) in 0.9% NaCl into the dorsal funiculus of the spinal cord at the level of the 13th thoracic vertebrate. Forty-eight hours after demyelination, mice received a single injection (1μl, 10^5^/μl) of GFP+ miPS-NPCs (n=6 /genotype) or mE-NPCs (n=6/genotype) or the same amount of medium (n=6/genotype) at the site of demyelination as previously described (Mozafari et al., 2015). All injections (LPC or cells) were performed at low speed (1μl/2min) using a stereotaxic frame equipped with a micromanipulator and a Hamilton seringe. For sacrifice, mice were perfused transcardially with a solution of 1X PBS and 4% paraformaldehyde. Nude mice (n=18) were sacrificed at 1, 2 wpg (n=3 for each cell type or medium, and for each time point). Two intact animals were sacrificed at the age of 8-9 weeks. Shiverer/Rag2^−/−^ mice were sacrificed (n=3 for each cell type and time point) at 6 and 10 wpg, 6 and 10 weeks wpm. Three intact animals were sacrificed at the age of 16 weeks (intact mice). After dissection, tissues were post-fixed in the same fixative for 1 hour and then processed in 20% sucrose in 1X PBS overnight. Spinal cords (~ 12 mm, including the lesion site at the center) were transversally cut in 3-5 pieces (~ 3mm each) and serially ordered in small plastic containers, embedded in cryomatrix (Shandon), frozen in cold Isopenthane at –60°C and stored at –20°C until use. Serial transverse sections were performed at 12μm with a cryostat (Leica) and collected on 3 series of 10 slides each (30-50 sections/slide) and used for immunohistochemistry. Our published data showed that the density of the grafted GFP+ cells was similar for both miPS-NPCs and mE-NPCs at 1wpg and 2wpg confirming the reproducibility of the lesion/graft paradigm (Mozafari et al., 2015).

One series of *Shi/Shi:Rag2^−/−^mice* (n=3) was also engrafted with human iPSC-OPC, sacrificed at 12 wpg and handled as above (Ehrlich et al., 2017).

### Immunohistochemistry

Demyelination was revealed using anti-MOG (mouse IgG1 hybridoma Clone C18C5, 1:50) antibody. Transplanted mouse cells were identified with an antibody directed against GFP (GFP-1020, Aves, chicken, 1:200). Differentiated oligodendrocytes were recognized using anti-CC1 (Millipore, OP80, IgG2b, 1:100) or anti-human NOGOA for grafted human OPC (Santa Cruz Biotechnology, sc-11030, goat, 1:50) antibodies. Oligodendroglial lineage cells were identified using anti-Olig2 (Millipore, AB 9610, rabbit, 1:400). Mouse astrocytes were revealed by anti-GFAP (Z0334, Dako, rabbit, 1:500) and anti-nestin (Millipore, MAB353, IgG1, 1:200).

Exogenous myelin was revealed using anti-MBP (Merck Millipore, AB980, rabbit, 1:100) and Neurofilaments were identified using anti-NF200 (N0142, Sigma, IgG1, 1:200). Oligodendrocyte Cx47 and astrocyte Cx43 were revealed respectively with anti-Cx47 (Invitrogen, 4A11A2, IgG1, 1:200) and anti-Cx43 (Sigma-Aldrich, C6219, rabbit, 1:50). For MBP and MOG staining, slices were pretreated with ethanol (10 min at room temperature). For connexin staining, sections were pre-treated with methanol (10 min, −20°C). Sections were incubated with the corresponding secondary antibodies from Jackson Immunoresearch Europe, or Alexa-conjugated antibodies (Invitrogen) containing Hoechst dye (1mg/ml). A Carl Zeiss microscope equipped with ApoTome.2 was used for tissue scanning, cell visualization and imaging.

### Quantification

#### Demyelination and panglial loss extension

The extent of demyelination (area with loss of MOG) and panglial loss (area with loss of Cx47 and Cx43 staining) were evaluated based on the percentage of each related immuno-depleted field out of the total area of the dorsal funiculus (MOG+) at 1wpg as previously described (Mozafari et al., 2015). Briefly, for each animal, six coronal serial sections with 120 μm intervals, were collected from the middle of the lesion site and quantified for different groups (n=3 per group) of lesioned animals grafted with miPS-NPCs and mE-NPCs or injected with medium.

#### Mature oligodendrocytes and Cx47 dynamic expression

CC1^+^ cells (GFP^−^ endogenous or GFP+ grafted from both cell sources) expressing or not the Cx47 were quantified on transverse sections at the lesion site to determine the total number of CC1^+^ cells or the proportions of total CC1^+^ mature oligodendrocytes (endogenous or exogenous) expressing Cx47. For each animal, six coronal serial sections at 120 μm intervals, were collected from the middle of the lesion site. The lesion area was defined by MOG and/or Cx47 depletion, and cell density was identified by Hoechst staining of all cell nuclei on adjacent sections. Sections were first scanned at 20X to define the limits of the lesion and level of maximal demyelination and magnified at 40X to quantify the amount of GFP cells expressing Cx47. Cell counts were expressed as the percentage of total CC1^+^, total CC1^+^/GFP^−^ or total CC1^+^/GFP^+^ cells in the lesion area.

#### Donor-derived remyelination

Exogenous myelin was visualized by MBP staining on transverse sections collected from transplanted *Shi/Shi:Rag2^−/−^* (6 and 10 wpg) mice. At the lesion core, the percentage of NF^+^/MBP^+^ axons over total number NF^+^ axons was quantified from three different fields of 1000 μm^2^ each per coronal section (on confocal images) and for three sections apart of 120 μm intervals (total number of 9 different fields per animal) for all the grafted *Shi/Shi:Rag2^−/−^* mice at 6 wpg and 10 wpg according to our previously described method (Mozafari et al., 2015).

### Statistics

Data were analyzed using Two-way ANOVA followed by Tukey’s or Sidak’ multiple comparison tests or Mann Whitney test (for less than 2 groups). Non-normally distributed data were analyzed by the corresponding non-parametric tests. Statistical analysis was carried out using GraphPad Prism 8 software. Data were presented as mean ± standard error of the mean (SEM) for all statistical analysis. A *P* value of less than 0.05 was considered significant.

## Acknowledgments

This work was supported by the Progressive MS Alliance (PMSA, collaborative research network PA-1604-08492 (BRAVEinMS), and INSERM and ICM internal fundings to ABVE. SM was funded by European Committee for Treatment and Research in Multiple Sclerosis (ECTRIMS). This work was supported by the Italian Multiple Sclerosis Foundation (FISM), Project No. ‘Neural Stem Cells in MS’ (to GM).

## Supplementary materials

**Fig. S1.**
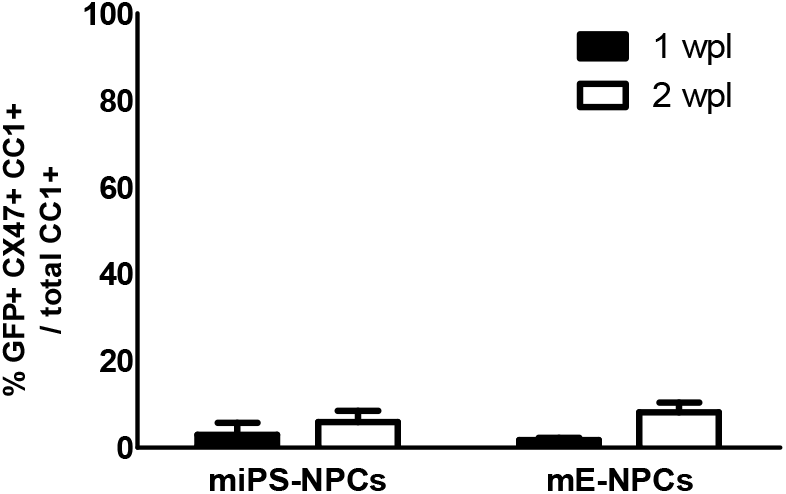
No difference in Cx47 expression over time between the two grafted cells types (miPSC-NPCs vs mE-NPCs).

**Fig. S2.**
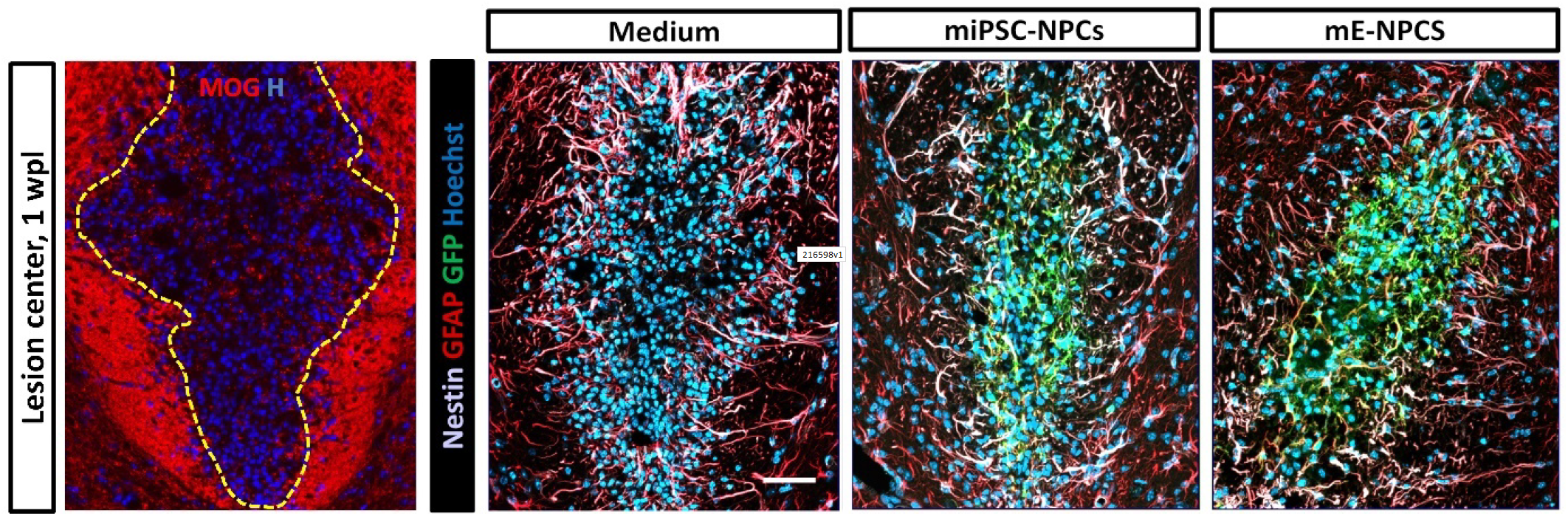
Astrocyte response to demyelination in medium, miPSC-NPC and mE-NPC injected mice revealed by GFAP (red) and nestin (gray) immunoreactivity 1 wpl. Some miPS-NPCs or mE-NPCs co-labelled with GFAP and nestin at the lesion site showing that few grafted cells differentiated into astrocytes.

